# Metabolic adjustments to winter severity in two geographically separated great tit (*Parus major*) populations

**DOI:** 10.1101/2023.08.01.551459

**Authors:** Cesare Pacioni, Andrey Bushuev, Marina Sentís, Anvar Kerimov, Elena Ivankina, Luc Lens, Diederik Strubbe

**Affiliations:** Terrestrial Ecology Unit, Ghent University, Ghent, Belgium; Department of Vertebrate Zoology, Faculty of Biology, M.V. Lomonosov Moscow State University, Moscow, Russia; S.N. Skadovsky Zvenigorod Biological Station, M.V. Lomonosov Moscow State University, Moscow, Russia

**Keywords:** Metabolic rates, Great tit, Aerobic capacity model, Basal metabolic rate, Summit metabolic rate

## Abstract

Understanding the potential limits placed on organisms by their ecophysiology is crucial for predicting their responses to varying environmental conditions. Studies to date have traditionally relied on between-species comparisons, however, recently, there has been a growing recognition of the importance of intraspecific variation in shaping an organism’s ecological and physiological responses. In this context, widely distributed resident bird species offer a well-suited study system to examine intraspecific geographical variation in ecophysiological traits. A main hypothesis for explaining avian thermoregulatory mechanisms is the aerobic capacity model, which posits a positive correlation between basal (BMR) and summit (M_sum_) metabolism, caused by the energetic maintenance costs associated with increased muscle mass for shivering thermogenesis and enhanced investment in digestive organs for food processing. Most evidence for this hypothesis, however, comes from interspecific comparisons only, and the ecophysiological underpinnings of avian thermoregulatory capacities hence remain controversial. Here, we focus on great tits (*Parus major*), measuring winter BMR and M_sum_ in two populations from different climates, a maritime-temperate (Gontrode, Belgium) and a continental (Zvenigorod, Russia) one. We test for the presence of intraspecific geographical variation in metabolic rates and assess the predictions following the aerobic capacity model. We found that metabolic rates differed between populations, whereby the birds from the maritime-temperate climate (Gontrode) showed higher (whole-body and mass-independent) BMR whereas conversely, great tits from Zvenigorod showed higher levels of both (whole-body and mass-independent) M_sum_. Within each population, our data did not fully support the aerobic capacity model’s predictions. We argue that the decoupling of BMR and M_sum_ observed may be caused by different selective forces acting on these metabolic rates, with birds from the continental-climate Zvenigorod population facing the need to conserve energy for surviving long winter nights (by keeping their BMR at low levels) while simultaneously being able to generate more heat (i.e., a high M_sum_) to withstand cold spells. We argue that the coupling or uncoupling of basal and maximum metabolic rates at the intraspecific level is likely influenced by different selective pressures that shape local adaptations in response to different climate regimes.

## Introduction

A key question in ecology is how species adjust their physiology to cope with different environmental conditions, as a better understanding of the underlying processes may allow ecologists to better predict how organisms will respond to changes in their environment (Bozinovic and Pörtner, 2015; Herrando-Pérez et al., 2023). This is particularly important for endotherms, as they need to maintain their core body temperature within a relatively narrow range. While endothermy allows animals to be active over a wide range of ambient temperatures, it comes at a potentially high energetic cost and places additional demands on their physiology, such as the need for efficient respiratory and cardiovascular systems and the ability to process food quickly and efficiently (Boyles et al., 2011; Kronfeld-Schor and Dayan, 2013). Studies attempting to unravel these physiological adjustments have relied heavily on interspecific comparisons, considering a given species as a homogenous physiological unit and assuming that conspecific populations have comparable responses (Reed et al., 2011; Thomas et al., 2004). However, different populations of a single species are likely to exhibit varying adjustments and different thermal tolerances to local environmental conditions (Cavieres and Sabat, 2008; Furness, 2003; Wikelski et al., 2003), as it is unlikely that a single phenotype will be the best fit for all conditions. This may be particularly true for populations that experience contrasting climatic conditions across their distribution range (Cavieres and Sabat, 2008; Root, 1988). Indeed, previous studies have shown that within-species variability in ecophysiological traits (e.g., metabolic rates) can be high, and by comparing individuals within a species, intraspecific studies can be used to test predictions derived from between-species comparisons and to identify factors beyond those revealed by interspecific studies (Cruz-Neto & Bozinovic, 2004).

Widely distributed resident bird species with geographical distributions spanning a range of climatic zones therefore provide an interesting study system for assessing intraspecific geographical variation in ecophysiological traits related to thermoregulation and providing functional explanations for it. The wide distribution suggests a role for local adaptation and/or phenotypic flexibility for population persistence in different climates (Kendeigh and Blem, 1974; Piersma and Drent, 2003). For instance, while interspecific studies have shown that differences in metabolic rates between species are largely due to differences in body mass, reflecting either local adaptation or phenotypic plasticity, functional explanations for this scaling remain controversial (Giancarli et al., 2023; White et al., 2019). In line with the ‘food-habitat’ hypothesis (Thompson, 2019), interspecific studies often document strong correlations between diet and metabolism, such as the finding that species feeding on a combination of insects with seeds and fruits typically have higher metabolic rates (McNab, 2009). However, intraspecific comparisons typically show mixed support for this hypothesis (Mckechnie & Swanson, 2010), suggesting that individual-level variability in factors such as enzymatic plasticity and the use of energy-saving mechanisms such as facultative torpor must be taken into account to understand the functional significance of such correlations (Cruz-Neto and Bozinovic, 2004).

Failure to identify, quantify, and explain intraspecific variation in thermoregulatory capacities can also lead to erroneous forecasts of range shifts due to climate change (Bozinovic et al., 2011; Pearman et al., 2010). To gain a comprehensive understanding of species’ responses to environmental changes, it is crucial to consider their thermoregulatory capacities (Boyles et al., 2011). Considerable uncertainty however remains regarding the extent to which local populations of a given species respond to weather and climate conditions across their range by adjusting their physiological characteristics. Birds from highly seasonal environments are known to be able to increase their cold tolerance in winter, which they achieve through physiological adjustments in, for example, body mass and metabolic rate (Swanson, 1990, 2010; Swanson and Olmstead, 1999; Swanson and Vézina, 2015). Several studies have documented that the magnitude of such thermogenic adjustments correlates with climate severity. For example, studies on house finch (*Carpodacus mexicanus*) populations living in different regions of North America, each with distinct climatic conditions, found that summit metabolism (M_sum_, which reflects maximal thermogenic capacity) was higher in colder areas (Dawson et al., 1983; O’Connor, 1996). However, Swanson (1993) compared M_sum_ in winter-acclimatized dark-eyed juncos (*Junco hyemalis*) from the cold winter climate of South Dakota (USA) and the milder winter climate of western Oregon (USA), but found that M_sum_ was similar between the two populations. More recently, Stager et al. (2021) studied dark-eyed juncos across North America and found that while thermogenic capacity was higher in colder areas, there was substantial variation among populations in the extent to which they adjusted their M_sum_ in response to climate.

Similar debates exist about the mechanisms underlying intraspecific variation in M_sum_. An important explanation for this comes from the aerobic capacity model for the evolution of endothermy (Bennett & Ruben, 1979), which assumes a positive correlation between basal and summit metabolism, e.g., due to energetic maintenance costs associated with increased muscle mass for shivering thermogenesis and/or increased investment in the gut and digestive organs to process enough food to fuel muscle thermogenesis (Mckechnie & Swanson, 2010). Between-species comparisons generally support the aerobic capacity model (Auer et al., 2017; Rezende et al., 2002). For example, Dutenhoffer and Swanson (1996) found a positive correlation between whole-body and mass-independent basal metabolic rate (BMR) and M_sum_ in ten different bird species from South Dakota, USA. Intraspecific studies, in contrast, provide inconclusive support for functional correlations between BMR and M_sum_ (Swanson et al., 2012). For example, Liknes and Swanson (1996) found that BMR and M_sum_ were positively correlated in winter and late summer for white-breasted nuthatches (*Sitta carolinensis*) and downy woodpeckers (*Picoides pubescens*) in South Dakota. However, at higher latitudes, winters are not only colder but are also characterized by shorter day lengths, limiting the time available for foraging. High maintenance costs (i.e., high BMR) during long nights may cause birds to exhaust their energy reserves, leading to selection against high winter BMR (Bozinovic and Sabat, 2010; Broggi et al., 2005). Indeed, O’Connor (1995) found that BMR was seasonally stable in house finches in Michigan, USA, whereas M_sum_ was higher in winter. Similarly, Le Pogam et al. (2020) demonstrated that snow buntings (*Plectrophenax nivalis*) increased their M_sum_ by about 25% during cold Canadian winters without a concomitant increase in BMR. Therefore, intraspecific studies examining variations in both metabolic rates have yielded inconsistent results, leaving uncertainties regarding the extent of variation within species.

Here, we study intraspecific geographic variation in ecophysiological traits hypothesized to underpin avian thermoregulation, using great tits (*Parus major*) as a case study. The great tit is one of the best studied bird species, breeding from approximately 10°S to 71°N, and remaining resident even at the northernmost limit of its breeding range (Cramp et al., 1993; Silverin, 1995). To this end, we measured the basal (BMR) and summit (M_sum_) metabolic rates in two populations of great tits, one living in a maritime-temperate climate characterized by mild winters (Belgium, Gontrode, Melle), and the other living in a continental climate characterized by long and cold winters (Russia, Zvenigorod Biological Station, Moscow Oblast). We predict (i) that individuals from the cold, continental population will be characterized by higher maximal thermogenic capacity (i.e., M_sum_) but lower maintenance costs (i.e., BMR) compared to those from the maritime temperate population, and that (ii) BMR and M_sum_ will be correlated within each population, following the aerobic capacity model.

## Materials and methods

### Study areas, trapping and maintenance

The research in Belgium (Gontrode, Melle) took place in the Aelmoeseneie forest (50.975°N, 3.802°E), covering an area of 28.5 hectares. The forest is a mixed deciduous forest surrounded by residential areas and agricultural fields. The fieldwork was carried out during the late winter (February 01 – March 11, 2022). Since autumn 2015, the forest has been equipped with 84 standard nest boxes specifically designed for great tits. The climate in this region is maritime-temperate, characterized by mild winters and constant rainfall throughout the year (corresponds to the *Cfb* subtype in the Köppen climate classification). To monitor the ambient temperature (T_a_) in the forest, 20 TMS-4 data loggers were deployed, positioned approximately 15 cm above the ground (Wild et al., 2019).

The research in Russia (Zvenigorod) took place at the Zvenigorod Biological Station (55.701°N, 36.723°E), affiliated with Lomonosov Moscow State University. The station territory encompasses two small settlements within the mixed forest of the Moskva River valley, as well as a predominantly spruce forest in the watershed area. The fieldwork was carried out during the late winter (January 22 – February 02, 2021; January 27 – February 23, 2022; March 04-17, 2023). Although the reserve spans a total territory of 715 hectares and comprises 540 nest boxes, the winter bird catching was carried out only near the feeder located at the center of one of the settlements. Additionally, a nighttime check was conducted only on the nearest nest boxes within a distance of 250 meters from the feeder. The climate of Zvenigorod region is temperate continental (corresponding to the *Dfb* subtype in the Köppen climate classification). To monitor the T_a_ in the forest, the DS1921G Thermochron iButton logger (Dallas Semiconductor) was used, positioned approximately 1.5 m above the ground near the feeder.

In both populations, bird capture consisted of nightly nest box checks and daily mist netting. The captured birds were transported to a nearby laboratory, where they were individually ringed for identification purposes. They were also assessed for age (1st winter or adult), sex (based on plumage characteristics), weighed to the nearest 0.1g (before and after each metabolic measurement), and placed in individual cages with access to food (mealworms and sunflower seeds) and water. Following the metabolic experiments, all birds were released at their original capture site. The study protocol in Belgium was approved by the Ethics Committee on Animal Experiments VIB/Faculty of Science of Ghent University (EC2020-063), and in Russia by the Bioethics Committee of Lomonosov Moscow State University (applications #120-a and #120-a-2 for the experimental procedures, and #10.2-hous. and 10.3-hous. for the short-term housing of birds).

In Gontrode (Table 1), we measured BMR in 40 individuals (19 males and 21 females, including 35 adults and 5 1st-winter). Of these, we also measured the M_sum_ of 36 individuals (16 males and 20 females, including 31 adults and 5 1st-winter). In Zvenigorod (Table 1) we measured BMR in 128 individuals (77 males and 51 females, including 51 adults and 77 1st-winter), and M_sum_ was measured in 20 individuals using flow-through respirometry (11 males and 9 females, including 6 adults and 14 1st-winter) and 35 individuals using closed-circuit respirometry (21 males and 13 females, including 14 adults and 20 1st-winter). Table 2 provides a summary of the T_a_ in both locations.

**Table 1.** Mean ± standard deviation of body mass (g), basal metabolic rate (BMR; ml O_2_/min), summit metabolic rate (M_sum_; ml O_2_/min) and metabolic expansibility (i.e., the ratio between M_sum_ and BMR; ME) in great tits (*Parus major*) from two different locations: Gontrode (Belgium) and Zvenigorod (Russia). The values for each location are further grouped by sex (female, F; males; M) and age (adult, ad; subadult; sad). The sample size *n* for each group is provided within brackets.

| Location | sex | age | Body mass (g) | BMR (ml O<sub>2</sub>/min) | M<sub>sum</sub> (ml O<sub>2</sub>/min) | ME (M<sub>sum</sub>/BMR) |
| --- | --- | --- | --- | --- | --- | --- |
| Gontrode | F | ad | 15.6 ± 1.1 (20) | 1.13 ± 0.12 (20) | 4.30 ± 0.37 (19) | 3.85 ± 0.48 (18) |
|  |  | sad | 17.0 (1) | 1.18 (1) | 5.33 (1) | 4.52 (1) |
|  | M | ad | 16.9 ± 1.6 (15) | 1.16 ± 0.16 (15) | 4.68 ± 0.21 (12) | 3.93 ± 0.47 (10) |
|  |  | sad | 18.1 ± 1.1 (4) | 1.35 ± 0.37 (4) | 5.19 ± 0.37 (4) | 4.03 ± 0.92 (4) |
|  | mean |  | 16.4 ± 1.5 | 1.17 ± 0.17 | 4.56 ± 0.45 | 3.92 ± 0.53 |
| Zvenigorod | F | ad | 17.1 ± 1.2 (21) | 1.05 ± 0.09 (21) | 6.80 ± 0.45 (3) | 6.47 ± 1.14 (3) |
|  |  | sad | 16.9 ± 0.9 (30) | 1.07 ± 0.06 (30) | 5.91 ± 0.51 (6) | 5.29 ± 0.88 (6) |
|  | M | ad | 18.1 ± 0.7 (30) | 1.10 ± 0.06 (30) | 6.41 ± 0.38 (3) | 6.03 ± 0.77 (3) |
|  |  | sad | 17.8 ± 0.8 (47) | 1.08 ± 0.07 (47) | 6.80 ± 0.62 (8) | 6.41 ± 0.40 (8) |
|  | mean |  | 17.6 ± 1.0 | 1.08 ± 0.07 | 6.48 ± 0.64 | 6.11 ± 0.81 |

**Table 2.**
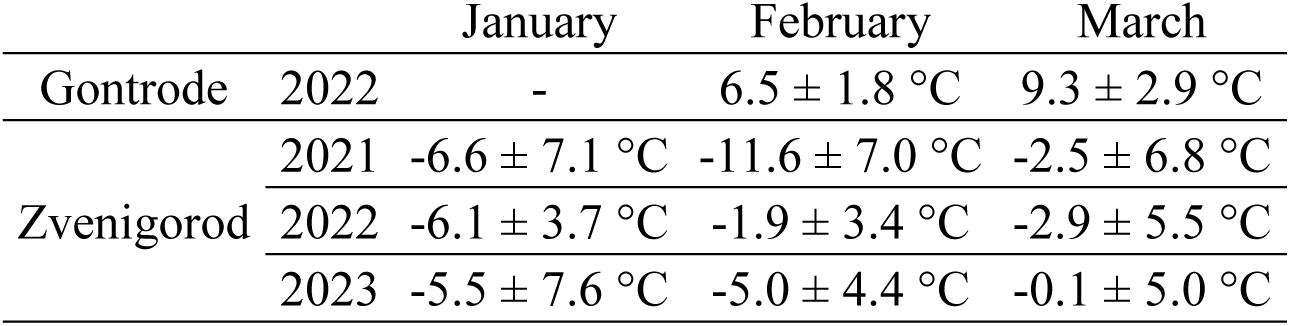
Mean ± standard deviation of ambient temperatures (T_a_) in both locations during the study periods.

### BMR measurements

The assessment of winter BMR in Gontrode was conducted at night using open flow-through respirometry (Lighton, 2018). To measure BMR, the oxygen consumption (VO_2_) of 40 individuals in a fasted state was monitored, following the methods described in Pacioni et al. (2023). Before and after the respirometry measurement, the body masses of the birds were taken to the nearest 0.1g. Each individual was then placed in a 1.1-liter plastic chamber within a darkened climate control unit (Combisteel R600). Ambient air was delivered by two pumps at a flow rate of 400 ml/min. The chambers were maintained at 25°C, which falls within the birds’ thermoneutral zone (Bech & Mariussen, 2022; Pacioni et al., 2023), determined according to the procedures outlined by van de Ven et al. (2013). The birds were measured in cycles, along with several baselines, with the timing and duration of measurements varying depending on the number of birds present during each session. On average, each bird underwent measurements for approximately 30 minutes per cycle, with three cycles conducted throughout the night. Following the metabolic measurements, the birds were returned to their cages and provided with water and food ad libitum. Additional details regarding calibration and the respirometry setup can be found in Pacioni et al. (2023).

The assessment of winter BMR in Zvenigorod was conducted similarly to BMR measurements in Gontrode. VO_2_ measurements were carried out throughout the night (from 14 h in late January to 11 h in mid-March) using flow-through respirometry. An 8-channel system enabled the measurement of VO_2_ in up to 7 birds (on average, 3.3 individuals). Outdoor air was pushed through columns containing silica gel. Subsequently, the dehumidified air was directed at an average flow rate of 430 ml/min into 1.25-liter polypropylene chambers housing the birds, which were maintained at a T_a_ of 26.5°C in thermostats. The air from the chambers was dried using a small chamber with 10-20 mesh Drierite® (W.A. Hammond Drierite Co. Ltd) and directed into the flowmeter of the FoxBox respirometer (Sable Systems). A subsample of the airflow, at a rate of 100 ml/min, was then directed to the O_2_ and CO_2_ analyzers in two FoxBoxes (the second one was used for control), which recorded gas concentrations and flow rates every 6 sec. The measurements of gas concentrations in the airflow from the chambers with birds and the reference chamber were alternated, with durations of 20-25 min for each bird and 5-10 min for the baseline measurement. The minimum VO_2_ (BMR) typically occurred around 3:30 am. Additional details regarding calibration and leakage testing can be found in Bushuev et al. (2021).

### M_sum_ measurements

During the day, individual measurements of winter M_sum_ were conducted in Gontrode on 36 individuals. The maximum cold-induced oxygen consumption (VO_2_) in a heliox atmosphere (79% helium, 21% oxygen) was used as the indicator for M_sum_, following the methodology outlined in Pacioni et al. (2023). The sliding cold exposure method of Swanson et al. (1996) was used. Prior to and after the trials, the body mass of each bird was recorded with an accuracy of 0.1g. Subsequently, the birds were placed in a 0.9-liter metal chamber. The chamber, along with the bird inside, was positioned within the same climate control unit used for BMR measurements, with an initial temperature setting of 10°C. The climate control unit was supplied with flowing heliox gas a few minutes prior to the trial, allowing the bird to acclimate. The heliox gas was pumped into the chamber at a flow rate of approximately 812 ml/min. Each M_sum_ trial began with a 7-minute baseline measurement using heliox, ensuring complete replacement of the air in the metabolic chambers before data recording commenced. After the baseline period, the experimental channel was activated. Following the removal of the bird from the chamber, baseline values were once again recorded for a minimum duration of 5 minutes. The trial was stopped when a steady decline in VO_2_ was observed for several minutes. M_sum_ was considered reached when the body temperature of a bird after a trial was 38°C (Cooper & Gessaman, 2005), measured by inserting a thermocouple (5SC-TT-TI-36-2M; Omega) coated with vaseline into the cloaca. The bird was then placed in a warm room with access to water and food. Additional details regarding calibration and the respirometry setup can be found in Pacioni et al. (2023).

In Zvenigorod, M_sum_ was estimated during the day using both flow-through respirometry (utilizing the same FoxBox system as for BMR measurements; n = 20) and closed-circuit respirometry (n = 35). The design of the flow-through respirometry setup was generally similar to that used in Gontrode, but the static cold exposure method described by Swanson et al. (1996) was employed. Before the start of the experiment, the birds were placed inside a 1.3-liter metal chamber with its interior painted black. Subsequently, the chamber was flushed with heliox until the volume of gas passing through the chamber reached six times its volume. The flow through the chamber was then reduced to 1.5 l/min, and a 5-minute reference measurement of concentrations of O_2_ and CO_2_ (baselining) was initiated. During this period, the chamber was carefully placed inside an Alpicool C40 refrigerator (avg. T_a_ = −10°C) filled with a mixture of propylene glycol, propanol, and water. The heliox flow from the gas cylinder passed through the FoxBox mass flow meter and then entered the chamber with the bird through a 20-meter-long tube, which was also located inside the refrigerator to allow the gas mixture to cool down. The experimental channel was activated every 20-25 min, alternating with 5-min baseline periods. After the chamber, the flow was subsampled at a rate of 200 ml/min. The subsampled gas was then passed through a 20 ml column containing Drierite® 10-20 mesh absorbent to remove water. Subsequently, the air passed through two FoxBoxes (the second one was used for control), which recorded O_2_ and CO_2_ concentrations every second.

To measure M_sum_ in a closed system, a respirometer constructed by D.V. Petrovski (Institute of Cytology and Genetics, Russian Academy of Sciences) was utilized. The bird, housed in a metal mesh cage, along with a permeable container containing CO_2_ and water absorbent (KOH granules), was placed inside a cylindrical steel chamber. This 1.2-liter chamber was located in WAECO TropiCool TC-35FL-AC thermoelectric refrigerator filled with antifreeze liquid (see above) at T_a_ = −1°C. Subsequently, the chamber was purged with heliox six times its volume. During the VO_2_ measurement, the heliox inside the airtight chamber with the bird was mixed using an internal membrane pump for a duration of 50 seconds. Following this, the decreased pressure within the chamber was balanced with atmospheric pressure by briefly opening a valve connecting the chamber to an oxygen pillow with pure O_2_ for 5 seconds. Another 5 seconds were dedicated to gas mixture mixing, so the cycle repeated every minute. The decrease in partial pressure of O_2_ in the chamber with the bird during each cycle was measured using an internal pressure gauge and converted to VO_2_/min. Further details can be found in Moshkin et al. (2002) and Vasilieva et al. (2020). Similar to Gontrode, a notable decrease in VO_2_ within a few minutes served as an indication of hypothermia. To measure cloacal temperature, we used a K-type thermocouple coated with vaseline and connected it to a calibrated Testo 175 T3 thermologger.

### Respirometry and data analysis

For the Gontrode data, BMR (ml O_2_/min) and M_sum_ (ml O_2_/min) were extracted using the ExpeData software provided by Sable Systems. The calculations for BMR, TNZ, and M_sum_ were performed using equation 9.7 from Lighton (2008). The lowest stable section of the curve, averaged over 5 minutes, was utilized to estimate BMR and TNZ throughout the entire night, while the highest 5-minute average VO_2_ during the test period was used to estimate M_sum_. To ensure accuracy, all data were adjusted for drift in O_2_, CO_2_, and H_2_O baselines utilizing the Drift Correction function available in ExpeData.

For the Zvenigorod data on BMR measurements, the calculation of VO_2_ from fractional concentrations of O_2_ and CO_2_ was performed using the equation from Bushuev et al. (2018). However, for the M_sum_ trials in the flow-through respirometer, the flow rate was estimated prior to the metabolic chamber. Therefore, the calculation of VO_2_ for these trials was conducted using a different equation, specifically equation 9.7 from Lighton (2008). In this equation, %H_2_O was set to zero, as water was eliminated using a chemical dryer. To estimate BMR and M_sum_, the minimum and maximum running average VO_2_ values over a 5-minute period were used, respectively. To account for drift in the O_2_ and CO_2_ baselines, linear correction was applied using two adjacent baselines: one before and one after the VO_2_ measurement. In closed-circuit respirometry, the VO_2_ readings were adjusted to standard conditions (STP) and then M_sum_ was calculated using the 5-minute maximum running average.

### Statistical analysis

Linear regression models with a Gaussian error distribution were used to test whether body mass, BMR, M_sum_, and metabolic expansibility (i.e., the ratio between M_sum_ and BMR; ME) differed between the two populations, specifying body mass, BMR, M_sum_, and, ME as dependent variables while adding location, sex, and age as covariates, with 2-way and 3-way interactions. Here, we followed Swanson et al. (1996), and considered M_sum_ values obtained from the sliding cold exposure method and the static cold exposure method to be comparable. Models were first run using whole-body metabolic rates, and then also using mass-independent metabolic rates. Mass-independent metabolic rates were considered as the residuals of regressions of (log) BMR, and (log) M_sum_ on (log) body mass (after the measurements). Prior to investigating potential intraspecific geographic variation in BMR, we assessed whether BMR and M_sum_ from Zvenigorod differed among the three years (2021, 2022, and 2023). Our analysis did not find any significant differences (p>0.1), hence we decided to consider BMR and M_sum_ measurements from three years as a single dataset for further analysis.

Regarding the assumption of the aerobic capacity model, similar linear models were used to test for positive correlations between body mass and metabolic rates, and between BMR and M_sum_. To avoid collinearity issues when assessing the relationship between mass-independent BMR and mass-independent M_sum_ we first calculated the residuals from the regression of (log) BMR and (log) M_sum_ on (log) body mass and then used the residual values of M_sum_ as the dependent variable and the residual value of BMR as the explanatory variable (Downs et al., 2013).

For all models, we used a backward stepwise procedure to eliminate non-significant interactions and variables. Post-hoc comparisons between species and seasons were performed with the emmeans function in the ‘emmeans’ package (Lenth, 2022). We used interquartile ranges as a criterion to identify outliers by using the quantile function. Then, we use the subset function to eliminate outliers. For all models, the normality of residuals was tested and verified (i.e., Shapiro-Wilk W > 0.9), and the significance level was set at p ≤ 0.05. Body mass, BMR, M_sum_, and ME were log-transformed before all analyses. Statistical analysis was performed using R v. 4.2.2 software (R Core Team, 2022).

## Results

### Variation in body mass and metabolic rate

We found a positive correlation between body mass and whole-body BMR, and between body mass and whole-body M_sum_ in both populations (BMR: Gontrode: R=0.48, p<0.01; Zvenigorod: R=0.58, p<0.0001; M_sum_: Gontrode: R=0.49, p<0.001; Zvenigorod: R=0.45, p<0.05). Individuals from Zvenigorod were significantly heavier (either considering the body mass before and after the metabolic measurements; p<0.05) than those from Gontrode (Gontrode: 16.4 ± 1.5g; Zvenigorod: 17.6 ± 1.0g; Table 1). Males were significantly (p<0.0001) heavier than females between and within populations, while adults and subadults did not differ in body mass between and within populations (p>0.05). Individuals from Gontrode had a significantly higher BMR than those from Zvenigorod, regardless of whether the analysis was conducted on whole-body (about two-fold higher, p<0.0001) or mass-independent (about four-fold higher, p<0.0001) metabolic rates. Males had a significantly higher whole-body BMR than females (p<0.05) between and within locations, due to their larger body mass, as mass-independent BMR did not differ between the sexes (p>0.1). Birds from Zvenigorod measured using the same open-flow respirometry set-up as used in Gontrode had a significantly higher M_sum_ (an increase of more than 30% (p<0.0001) for whole-body and almost two times higher (p<0.05) for mass-independent) than those from Gontrode. Great tits from Gontrode had a ME of 3.92 ± 0.53, whereas great tits from Zvenigorod had a whole-body ME of 5.83 ± 1.52. Whole-body ME from Zvenigorod was significantly higher (about 50%, p<0.0001) compared to birds from Gontrode (Figure 1). When considering the Zvenigorod birds whose M_sum_ was measured using closed-flow respirometry, differences between Gontrode and Zvenigorod great tits were even more outspoken (e.g., 1.5 times higher mass-independent M_sum_; see Supplementary file 1 for details). In the remainder of the manuscript, we therefore focus on the more conservative and comparable open-flow metabolic rates.

**Figure 1.**
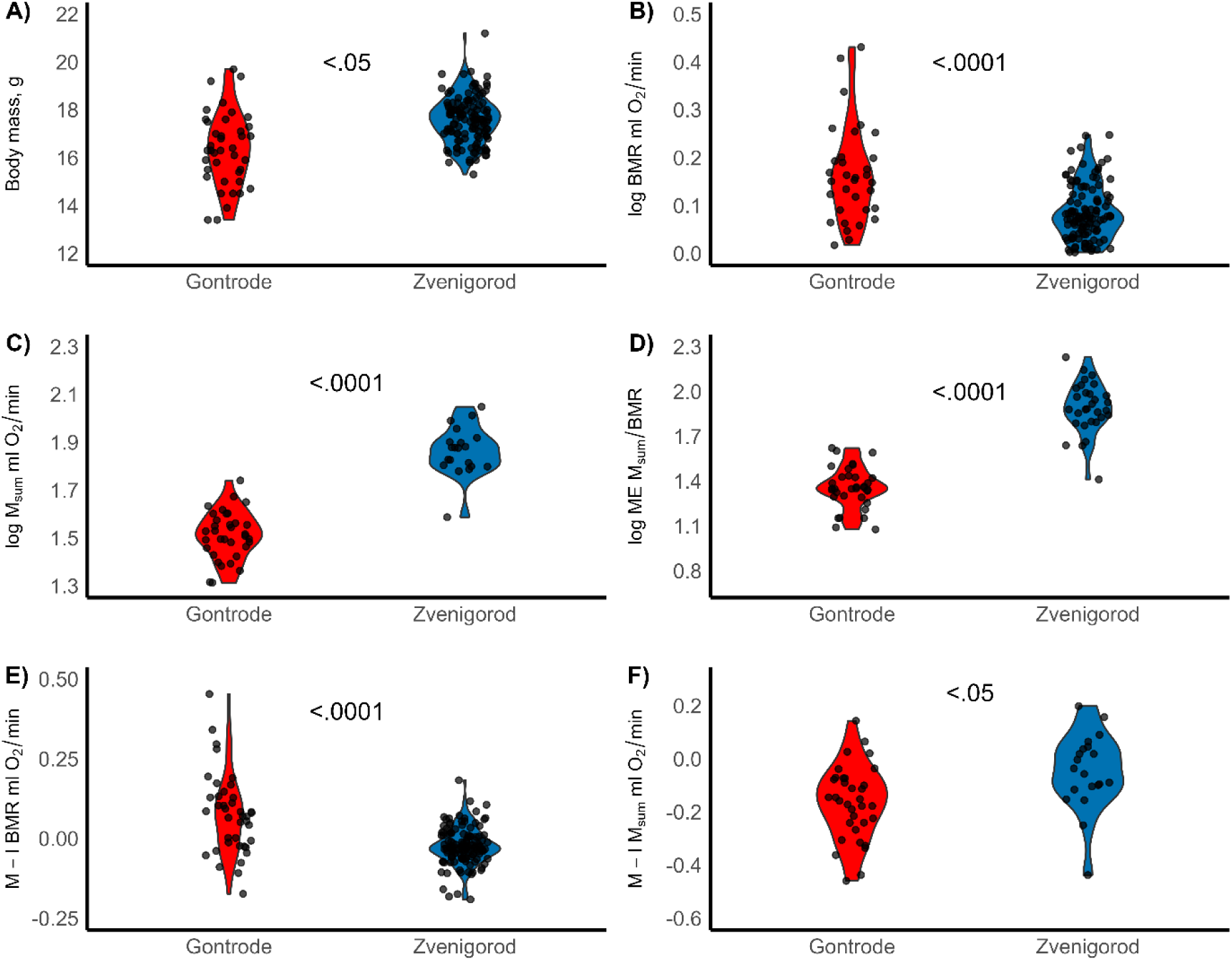
Violin plots of A) body mass (g), B) whole-body basal metabolic rate (BMR; ml O_2_/min), C) whole-body summit metabolic rate (M_sum_; ml O_2_/min), D) whole-body ME (M_sum_/BMR), E) mass-independent (M-I) BMR and F) mass-independent (M-I) M_sum_ in great tits (*Parus major*) from two different locations: Gontrode (Belgium, red) and Zvenigorod (Russia, blue). The p-value indicating the statistical significance of the differences is displayed between the two corresponding box plots.

### Aerobic capacity model of endothermy predictions

Whole-body BMR and M_sum_ were also positively correlated in individuals from Gontrode (R=0.41, p<0.05), while mass-independent metabolic rates were not (p>0.1). In Zvenigorod, no significant correlations were found between BMR (whole-body or mass-independent) and M_sum_ (Figure 2). Details of the results of all statistical analyses are available in Supplementary file 1.

**Figure 2.**
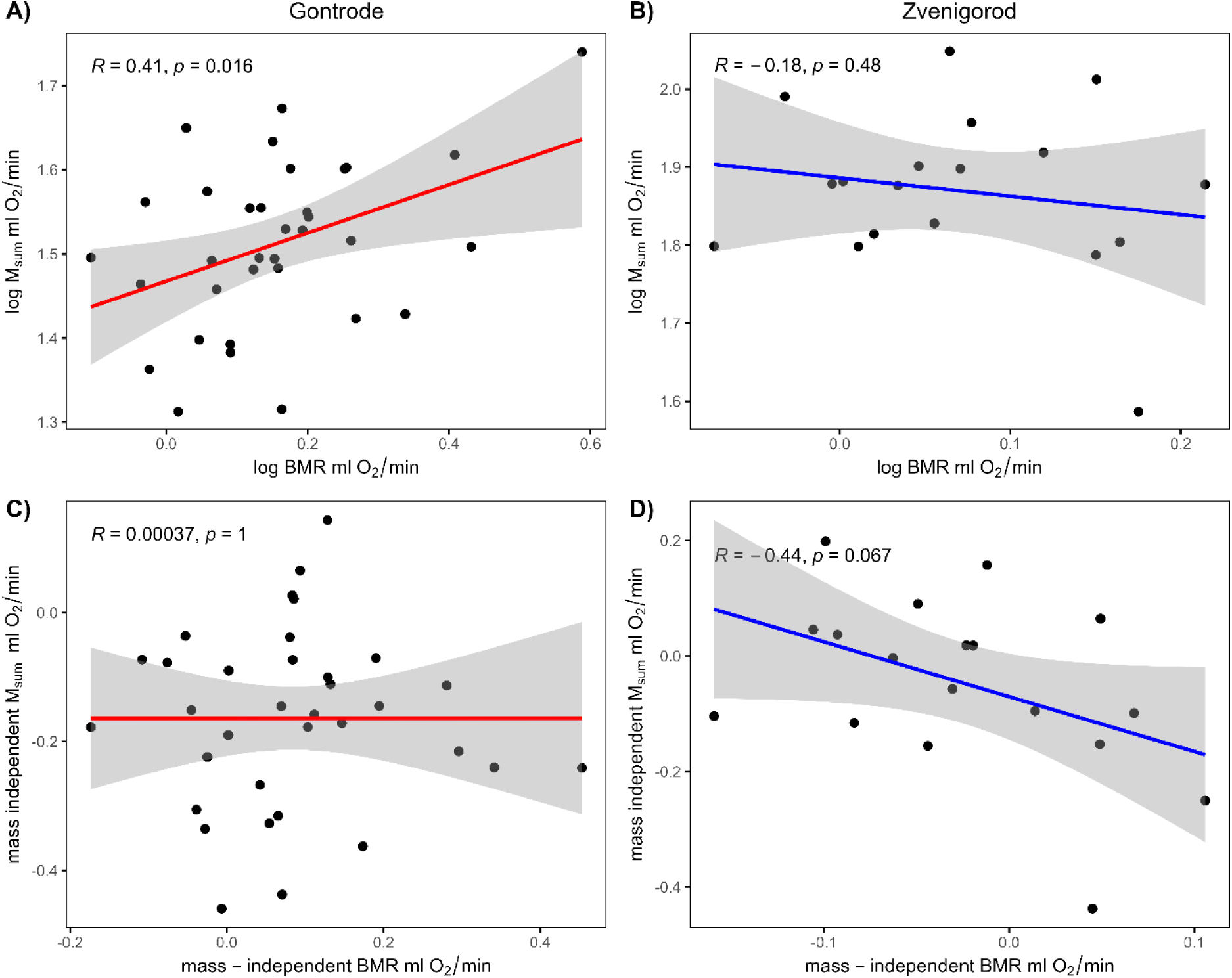
Relationships between (log) BMR and (log) M_sum_ in Gontrode, Belgium (A) and Zvenigorod, Russia (B) and between mass-independent BMR and mass-independent M_sum_ in Gontrode (C) and Zvenigorod (D).

## Discussion

Here, we test for intraspecific variation in ecophysiological traits related to thermoregulation using great tits living in two geographically and climatically separate locations that experience different winter conditions. While several studies have documented that metabolic rates can vary between geographical areas, this study is, to our knowledge, the first to tests for concurrent variation in basal (BMR) and summit (M_sum_) metabolic rates. Observed differences between populations suggest that avian basal and summit metabolic rates may vary independently in response to environmental influences, as great tits from the colder site (Zvenigorod, Russia) had significantly higher thermogenic capacity (i.e., M_sum_) than those from the warmer site (Gontrode, Belgium), but a lower basal metabolic rate (BMR). Contrary to the prediction of the aerobic capacity model, we find only weak support for a functional relationship between these metabolic rates at the individual level as (mass-independent) BMR and M_sum_ were uncorrelated.

During winter, continental high-latitude areas are characterized by low temperatures, limited food resources, shorter foraging periods, and extended fasting periods during the long nights. Physiological adaptations have long been hypothesized to be key to enduring such challenging environments (Swanson and Olmstead, 1999). Indeed, more recently, Petit et al. (2017), for example, found that winter survival of black-capped chickadees (*Poecile atricapillus*) in Quebec, Canada, was positively associated with M_sum_. However, the relationship between winter survival and M_sum_ was not linear, but exhibited a threshold function, and many individuals had M_sum_ values significantly higher than required for survival. Petit et al. (2017) suggested that this increase in M_sum_ may be due to selection for increased muscle mass to power the sustained foraging flights required to gather sufficient food in the generally resource-poor winter habitat, with the thermogenic effects providing a secondary benefit. Such an explanation for high M_sum_ is less likely here, as higher muscle mass should correspond to higher overall energetic maintenance costs, which was not confirmed by our data. Moreover, birds from the colder Zvenigorod site were characterized by lower whole-body BMR compared to birds from Gontrode, despite having higher body mass. This suggests that the differences in mass between the populations are not due to a higher muscle mass, but rather that the Zvenigorod birds may have a greater amount of metabolically (relatively) inactive fat mass (Scott and Evans, 1992). Indeed, the accumulation of fat reserves during winter, known as winter fattening, is a well-known mechanism observed in small passerines to survive colder climates and longer nights (King and Farner, 1966; Pravosudov and Grubb, 1997).

The Zvenigorod population thus has a higher metabolic expansibility (whole-body M_sum_/BMR; ME: ~6 times the BMR compared to ~4 times for the Gontrode population), which is commonly used as an indicator of the organism’s ability to produce heat for a given level of metabolic maintenance cost (Arens and Cooper, 2005; Cooper and Swanson, 1994). As we found no support for the aerobic capacity model of endothermy, which postulates a positive correlation between minimum (BMR) and maximum (M_sum_) aerobic metabolic rates (Bennett and Ruben, 1979), it remains unclear what mechanisms underlie the summit metabolic capacity of our study birds. The aerobic model has been supported primarily by interspecific studies (Dutenhoffer and Swanson, 1996; Rezende et al., 2002), but these relationships often disappear at the individual level, at least after accounting for variation in body mass (Swanson et al., 2012; Vézina et al., 2006). The reasons for the differences between inter- and intraspecific studies of these phenotypic correlations are not yet fully understood. On the one hand, this lack of correlations may represent a statistical artefact associated with much lower levels of variation in body mass and metabolic rates in intraspecific studies (Swanson et al., 2017). On the other hand, Swanson et al. (2023) carried out a literature review investigating flexibility in BMR, M_sum_ and metabolic expansibility, and found that, in fact, for none of the six species for which data were available, higher levels of flexibility in M_sum_ or metabolic expansibility did not typically result in increased maintenance costs (i.e., BMR). This suggests that non-shivering thermogenesis mechanisms may also contribute significantly to thermoregulation in birds (Pani & Bal, 2022). Indeed, multiple adaptations at the cellular and biochemical level have been shown to affect an organism’s thermogenic capacity when exposed to changing temperatures (Milbergue et al., 2018; Swanson, 2010). For example, Nord et al. (2021) recently showed that resident great tits in western Scotland were able to upregulate mitochondrial respiration rate and mitochondrial volume in winter, thereby increasing thermogenic capacity at the subcellular level. Further intraspecific studies should consider measuring and accounting for the body composition of individual birds, in particular the relative contributions of fat and muscle mass to the observed body mass differences, as well as blood sampling to investigate the possible role of non-shivering thermogenesis.

Alternatively, the decoupling of BMR and M_sum_ often observed in intraspecific studies may be caused by different selective forces acting on these metabolic rates (Petit et al., 2013), with birds from the Zvenigorod population facing the need to conserve energy (by keeping their BMR at comparatively low levels) while simultaneously increasing their thermogenic capacity (by increasing their M_sum_). Swanson et al. (2017) argues that, from an energetic point of view, natural selection should generally favor reducing BMR to the lowest possible level under prevailing environmental or ecological demands, allowing energy to be allocated to other functions. Bozinovic and Sabat (2010) similarly suggested that in resource-poor habitats, organisms that can reduce their BMR will reduce their daily energy expenditure and hence food requirements, thereby increasing fitness through increased survival. Broggi et al. (2005) used a common garden experiment to investigate the physiological basis of interpopulation differences in BMR in Scandinavian great tits, and found that selection pressure for low BMR was particularly strong in more northerly populations, where the energetic costs of thermoregulation and activity are highest. More recently, a literature review by Stager et al. (2016) suggested that latitudinal trends in metabolic rate are primarily driven by a necessary balance between increased thermogenic capacity to cope with cold temperatures and pressure to reduce excess maintenance costs in cold and low food availability environments.

Our results are therefore consistent with the expectation that birds in resource-poor, cold environments will be characterized by low BMR but high M_sum_. In contrast, in Gontrode, temperatures are never very cold and birds had access to abundant and stable food resources throughout the winter because the forest is surrounded by a residential area with several gardens equipped with bird feeders (Dekeukeleire, 2021). This may explain their lower body mass compared to the Zvenigorod birds, as the rich and stable food may allow birds to adopt a lean body mass by trading off predation and starvation risk (Macleod et al., 2005). Ambient air temperatures at Gontrode in February and March averaged 6.5 ± 1.8 °C and 9.3 ± 2.9 °C respectively (Table 2), compared to a normal average temperature of about 4.2 °C for February, and about 7.1 °C for March (Pacioni et al., 2023). Due to these comparatively warm temperatures, relatively few great tits were found roosting in nest boxes, further suggesting that thermoregulatory demands and starvation risks were less severe for Gontrode birds, weakening selection for lower BMR in this population.

### Conclusions

In conclusion, great tits from the colder site (Zvenigorod, Russia) had significantly higher thermogenic capacity (i.e., M_sum_) than those from the warmer site (Gontrode, Belgium), but a lower basal metabolic rate (BMR). This contradicts the aerobic capacity model, showing that avian basal and summit metabolic rates may vary independently in response to environmental influences. The coupling or uncoupling of minimum and maximum metabolic rates at the intraspecific level may then be influenced by different selective pressures that shape local adaptations in response to different degrees of seasonality. Therefore, future intraspecific studies should consider the potential impact of local adaptations and selective pressures on different conspecific populations when testing the applicability of the aerobic capacity model. These findings would contribute to a better understanding of the adaptive strategies used by birds in different environments and highlight the importance of considering multiple factors when studying and comparing avian metabolic rates.

## Acknowledgements

We express our gratitude to ForNaLab (Gontrode) for giving us access to their facilities. We extend special thanks to Luc Willems for his invaluable support and to Dries Landuyt for providing us with the temperature data from the dataloggers (Gontrode). We are grateful to the administration of the Zvenigorod biological station for providing the necessary conditions for the study. Furthermore, we would like to acknowledge the valuable assistance provided by bachelor student Hanne Danneels during the data collection process. This study acknowledges funding by FWO-Vlaanderen (project G0E4320N), by Methusalem Project 01M00221 (Ghent University) awarded to Frederick Verbruggen, Luc Lens, and An Martel, and by the Russian Science Foundation (project 20-44-01005). Marina Sentís acknowledges the support of FWO-Vlaanderen (project 11E1623N).

## Author contributions

Cesare Pacioni: Conceptualization, Methodology, Validation, Formal analysis, Investigation, Data curation, Writing – original draft, Visualization. Andrey Bushuev: Conceptualization, Methodology, Validation, Formal analysis, Investigation, Data curation, Writing – review & editing. Marina Sentís: Conceptualization, Methodology, Investigation, Writing – review & editing. Anvar Kerimov: Conceptualization, Methodology, Validation, Formal analysis, Investigation, Data curation, Writing – review & editing. Elena Ivankina: Methodology, Investigation, Writing – review & editing. Luc Lens: Conceptualization, Validation, Writing – review & editing, Supervision. Diederik Strubbe: Conceptualization, Methodology, Validation, Investigation, Writing – review & editing, Visualization, Supervision.

## Data availability

Script and data are made available on Mendeley Data (DOI: 10.17632/t84ntxxcbn.1)

